# Description of *Klebsiella spallanzanii* sp. nov. and of *Klebsiella pasteurii* sp. nov

**DOI:** 10.1101/703363

**Authors:** Cristina Merla, Carla Rodrigues, Virginie Passet, Marta Corbella, Harry A. Thorpe, Teemu V.S. Kallonen, Zhiyong Zong, Piero Marone, Claudio Bandi, Davide Sassera, Jukka Corander, Edward J. Feil, Sylvain Brisse

## Abstract

*Klebsiella oxytoca* causes opportunistic human infections and post-antibiotic haemorrhagic diarrhoea. This *Enterobacteriaceae* species is genetically heterogeneous and is currently subdivided into seven phylogroups (Ko1 to Ko4, Ko6 to Ko8). Here we investigated the taxonomic status of phylogroups Ko3 and Ko4. Genomic sequence-based phylogenetic analyses demonstrate that Ko3 and Ko4 formed well-defined sequence clusters related to, but distinct from, *Klebsiella michiganensis* (Ko1), *Klebsiella oxytoca* (Ko2), *K. huaxiensis* (Ko8) and *K. grimontii* (Ko6). The average nucleotide identity of Ko3 and Ko4 were 90.7% with *K. huaxiensis* and 95.5% with *K. grimontii*, respectively. In addition, three strains of *K. huaxiensis*, a species so far described based on a single strain from a urinary tract infection patient in China, were isolated from cattle and human faeces. Biochemical and MALDI-ToF mass spectrometry analysis allowed differentiating Ko3, Ko4 and Ko8 from the other *K. oxytoca* species. Based on these results, we propose the names *Klebsiella spallanzanii* for the Ko3 phylogroup, with SPARK_775_C1^T^ (CIP 111695^T^, DSM 109531^T^) as type strain, and *Klebsiella pasteurii* for Ko4, with SPARK_836_C1^T^ (CIP 111696^T^, DSM 109530^T^) as type strain. Strains of *K. spallanzanii* were isolated from human urine, cow faeces and farm surfaces, while strains of *K. pasteurii* were found in faecal carriage from humans, cows and turtles.

**Accession numbers:** The nucleotide sequences generated in this study were deposited in ENA and are available through the INSDC databases under accession numbers MN091365 (SB6411^T^ = SPARK775C1^T^), MN091366 (SB6412 ^T^ = SPARK836C1^T^) and MN104661 to MN104677 (16S rRNA), MN076606 to MN076643 (*gyrA* and *rpoB*), and MN030558 to MN030567 (*bla*_OXY_). Complete genomic sequences were submitted to European Nucleotide Archive under the BioProject number PRJEB15325.

**Abbreviations:** ANI, average nucleotide identity; HCCA, a-cyano-4-hydroxycinnamic acid; isDDH, *in silico* DNA-DNA hybridization; SCAI, Simmons citrate agar with inositol; MALDI57 ToF MS: Matrix-assisted laser desorption/ionization time of flight mass spectrometry

## Introduction

The genus *Klebsiella*, a member of the *Enterobacteriaceae* family, includes Gram-negative, non-motile (except *K. aerogenes*) and non-spore-forming capsulated bacteria. Bacteria belonging to the genus *Klebsiella* are found in water, soil and plants, and as commensals in the gut of animals including humans (1,2,3). In humans, *Klebsiella* species are frequently associated with hospital-acquired infections and are increasingly multidrug-resistant (4). *Klebsiella oxytoca* is the second most common *Klebsiella* species causing disease in humans, after *K. pneumoniae* (5). *K. oxytoca* carries a chromosomally encoded β-lactamase gene (*bla*_OXY_) that confers resistance to amino- and carboxypenicillins (6). This gene was shown to have diversified in parallel to housekeeping genes, and variants were classified into seven groups (*bla*_OXY-1_ to *bla*_OXY-7_) (7, 8, 9, 10). *K. oxytoca* phylogenetic lineages were named Ko1, Ko2, Ko3, Ko4, Ko6 and Ko7 reflecting which *bla*_OXY_ variant they carry; note that Ko5 was not defined, as isolates carrying *bla*_OXY-5_ represent a sublineage of Ko1 (9). Taxonomic work has shown that *K. oxytoca* (*sensu lato, i.e.*, as commonly identified in clinical microbiology laboratories) is in fact a complex of species, with *K. oxytoca* (*sensu stricto*) corresponding to phylogroup Ko2, *K. michiganensis* to Ko1 (11) and *K. grimontii* to Ko6 (12). The closely related *K. huaxiensis* (13) represents yet another phylogroup, which we here denominate as Ko8 and which carries *bla*_OXY-8_. Phylogroups Ko3, Ko4, Ko7 and *K. huaxiensis* were so far described only based on a single strain (9, 13), which has limited our ability to define their genotypic and phenotypic characteristics. While analysing a large number of *Klebsiella* strains from multiple human, animal and environmental sources in and around the Northern Italian town of Pavia, we identified 3 Ko3, 13 Ko4 and 3 *K. huaxiensis* strains. The aim of this work was to define the taxonomic status of *K. oxytoca* phylogroups Ko3 and Ko4 and provide identification biomarkers for all members of the *K. oxytoca* species complex.

## Material and methods

### Bacterial strains

Novel strains (3 Ko3, 13 Ko4 and 3 Ko8) were isolated through enrichment in Luria-Bertani broth supplemented with 10 μg/mL of amoxicillin, followed by isolation on Simmons citrate agar with 1% inositol (SCAI) medium (14) and re-isolation on MacConkey agar. Additional strains, including type and reference strains of each *K. oxytoca* phylogroup and the type strain of *K. pneumoniae* (15) were included in the study (**Table 1**). Strain SG271 (internal strain bank identifier, SB3356) and SG266 (SB3355) were included as reference strains for the phylogroups Ko3 and Ko4, respectively (9).

**Table 1.**
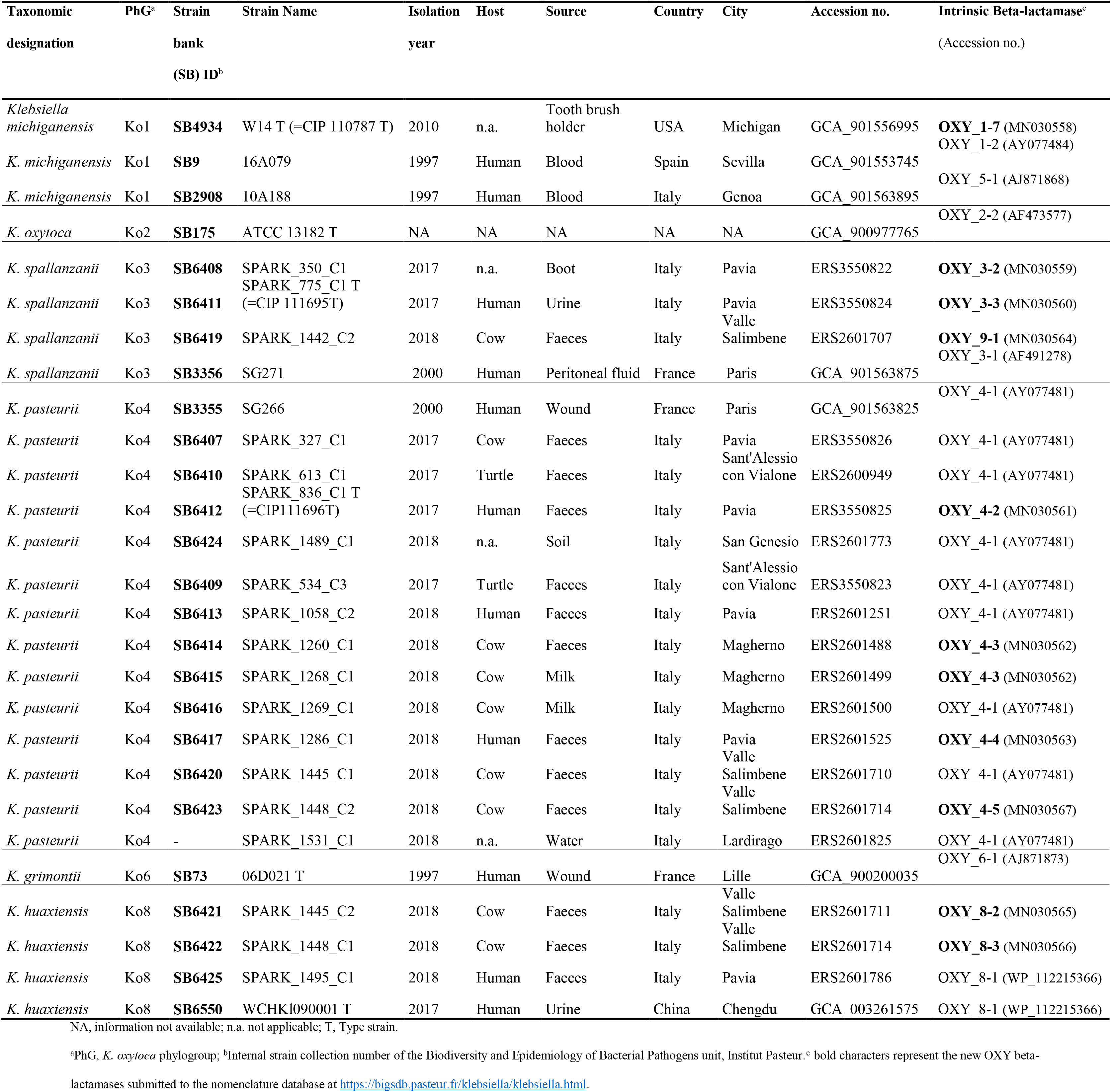
Strains included in the study, with provenance and genomic information

### Genome sequencing and analyses

Colonies from the novel strains grown on MacConkey agar were collected and resuspended in distilled water for DNA purification, which was performed using QIAsymphony automated instrument with the kit QIAsymphony DSP Virus/Pathogen following the manufacturer’s recommendation. DNA was stored at −20°C until sequencing on an Illumina HiSeq X Ten platform with a 2 x 150 nt paired-end protocol. Reads were assembled using SPAdes v3.11 and the assemblies were annotated using Prokka v1.12 (16). JSpeciesWS (17) was used to calculate the average nucleotide identity (ANI) using the BLAST algorithm (ANIb), whereas *in silico* DNA-DNA hybridization (isDDH) was performed through GGDC tool (http://ggdc.dsmz.de; formula 2) (18). Sequences of *gyrA* and *rpoB* genes were obtained from genome assemblies using BLASTN, while 16S rRNA gene sequences were obtained using Barrnap (https://github.com/tseemann/barrnap). The chromosomal *bla*_OXY_ sequences were also extracted, and the new amino-acid sequence variants were submitted to the Institut Pasteur MLST nomenclature database (https://bigsdb.pasteur.fr/klebsiella) for variant number attribution, and to NCBI for accession number attribution. 16S rRNA, g*yrA*, *rpoB* and *bla*_OXY_ beta-lactamase gene sequences were aligned using Muscle (19), concatenated (in the case of *rpoB* and *gyrB*) and phylogenetic relationships were assessed using MEGA v7.0 (20). Genetic distances were inferred using the neighbor-joining method with the Jukes-Cantor correction (21) in the case of nucleotide sequences or maximum-likelihood with Jones-Taylor-Thornton (JTT) (22) model in the case of the beta-lactamase protein sequences. The genome-based phylogenetic analysis was performed on the concatenation of 3,814 core genes defined using Roary v3.12 (23) with a blastP identity cut-off of 80% and presence in more than 90% of the isolates. *K. pneumoniae* ATCC 13883^T^ (GCA_000742135.1) was used as outgroup. An approximate maximum-likelihood phylogenetic tree was inferred using FastTree v2.1 (24).

### Biochemical and proteomic analyses

A representative subset of strains (n=30, 7 Ko1, 5 Ko2, 4 Ko3, 5 Ko4, 6 Ko6, 3 Ko8) of phylogroups of the *K. oxytoca* complex was subjected to API20E (BioMérieux) and to phenotype microarray characterization using plates PM1 and PM2 (Biolog, Hayward, CA, USA) in aerobic conditions as previously described by Blin and colleagues (25). The same subset of strains was also used to perform a MALDI-ToF mass spectrometry (MS) analysis following the protocol described by Rodrigues *et al.* (26). Briefly, cell extracts were spotted onto an MBT Biotarget 96 target plate, air dried and overlaid with 1 μL of a saturated α-cyano-4-hydroxycinnamic acid (HCCA). Mass spectra were acquired on a Microflex LT mass spectrometer (Bruker Daltonics, Bremen, Germany) using the default parameters, preprocessed (applying smoothing and baseline subtraction) with FlexAnalysis software, and then imported and analyzed in a dedicated BioNumerics v7.6 (Applied-Maths, Belgium) database.

## Results

The genome-based phylogenetic analysis based on the concatenation of 3,814 core genes (**Figure 1**) showed six distinct and highly supported branches. The thirteen Ko4 strains were clustered with Ko4 reference strain SG266 (SB3355) and this group was related to, but clearly distinct from, *K. grimontii* (Ko6). The three Ko3 strains (SPARK_350_C1, SPARK_775_C1 and SPARK_1442_C1) formed a well-defined cluster with Ko3 reference strain SG271 (SB3356, **Figure 1**), whereas the remaining three strains (SPARK_1445_C2, SPARK_1448_C1, SPARK_1495_C1) clustered with *K. huaxiensis*, which formed a distinct phylogroup that we here name Ko8. We therefore identified novel strains of these three phylogroups, which were each previously recognized based on a single strain. Furthermore, genome-based phylogeny revealed that Ko4 shares a common ancestor with *K. grimontii*, *K. michiganensis* and *K. oxytoca*, whereas Ko3 and *K. huaxiensis* share a common ancestor distinct from the Ko1/Ko4/Ko6 one (**Figure 1**).

**Figure 1.**
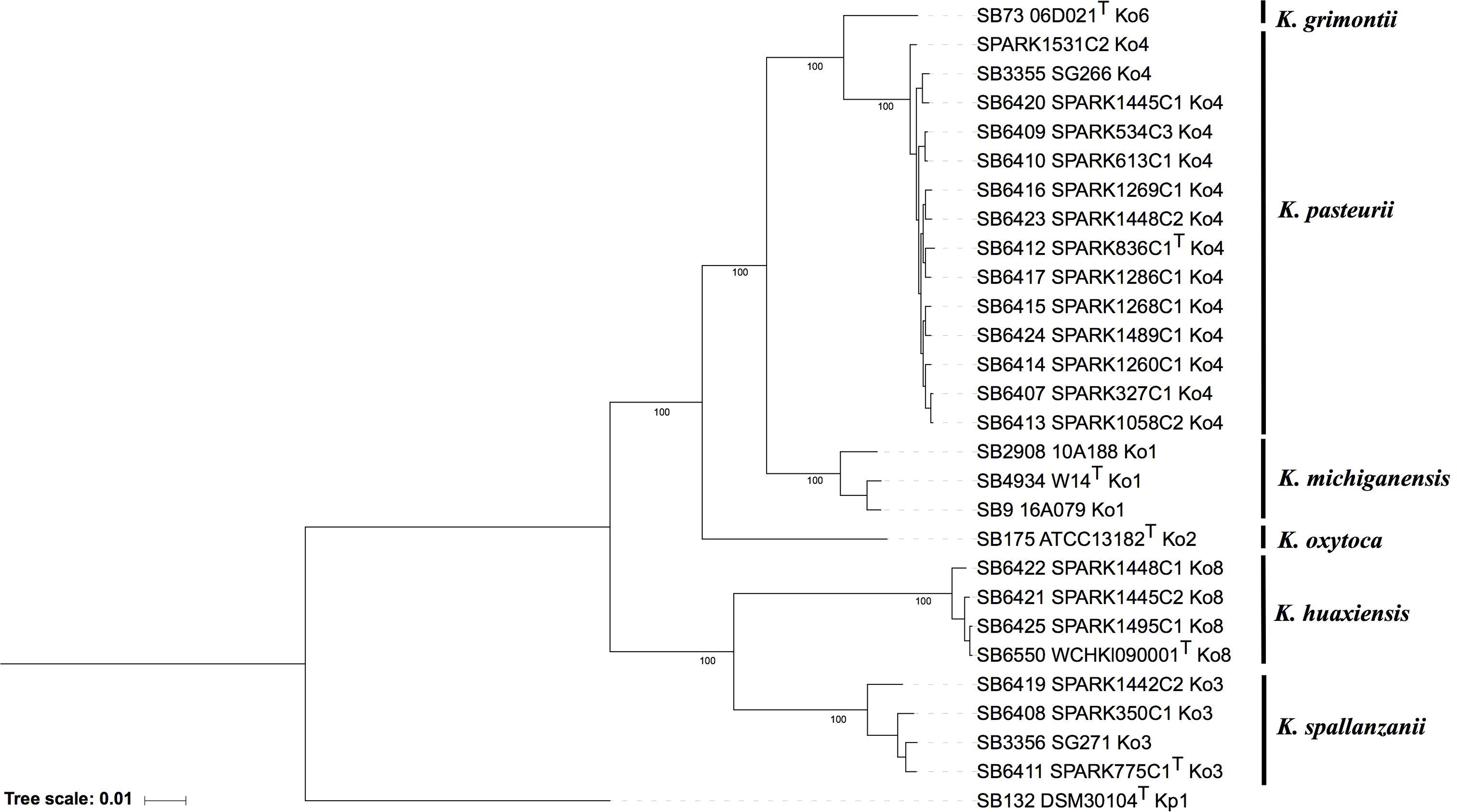
Maximum likelihood phylogenetic tree inferred based on the concatenated nucleotide sequence alignments of 3,814 core genes. The tree was rooted using *K. pneumoniae* DSM 30104^T^ (=ATCC 13883^T^). Taxonomic groups are indicated in front of the branches. Branch lengths represent the number of nucleotide substitutions per site (scale, 0.01 substitution per site). Bootstrap values are indicated at major nodes. Strain labels are given as Strain Bank ID (*e.g.*, SB73) followed by original strain name, followed by the phylogroup. A ‘T’ after the strain name indicates that the strain is the type strain of its taxon.

To determine how previously-used phylogenetic markers (7, 8, 9, 27) would group these novel strains, the sequences of internal portions of the housekeeping genes *gyrA* (383 nt) and *rpoB* (501 nt), as well as the *rrs* (1,454 nt) sequence coding for 16S rRNA, were extracted from genomic sequences and compared to previously characterized sequences of reference and type strains from the *K. oxytoca* complex (**Table 1**). The clustering of Ko4 strains and Ko3 strains was supported by phylogenetic analysis of combined *gyrA* and *rpoB* gene sequences (**Figure 2**), as well as by single gene phylogenies (**Figures S1 and S2**), showing that either gene used alone would allow reliable identification. The phylogeny of the chromosomal OXY beta-lactamase gene (**Figure S3**) was also in concordance with previous phylogenetic analyses. However, phylogroup Ko1 and Ko3 each harbored two different types of *bla*_OXY_, coding for OXY-1/OXY-5 and OXY-3/OXY-9, respectively (**Figure S3**). As previously reported (12, 28, 29), the phylogeny based on the *rrs* gene was not reliable for species or phylogroup identification (type strain sequences were > 97.8% similar), with only a few informative variable sites (**Figure S4**).

**Figure 2.**
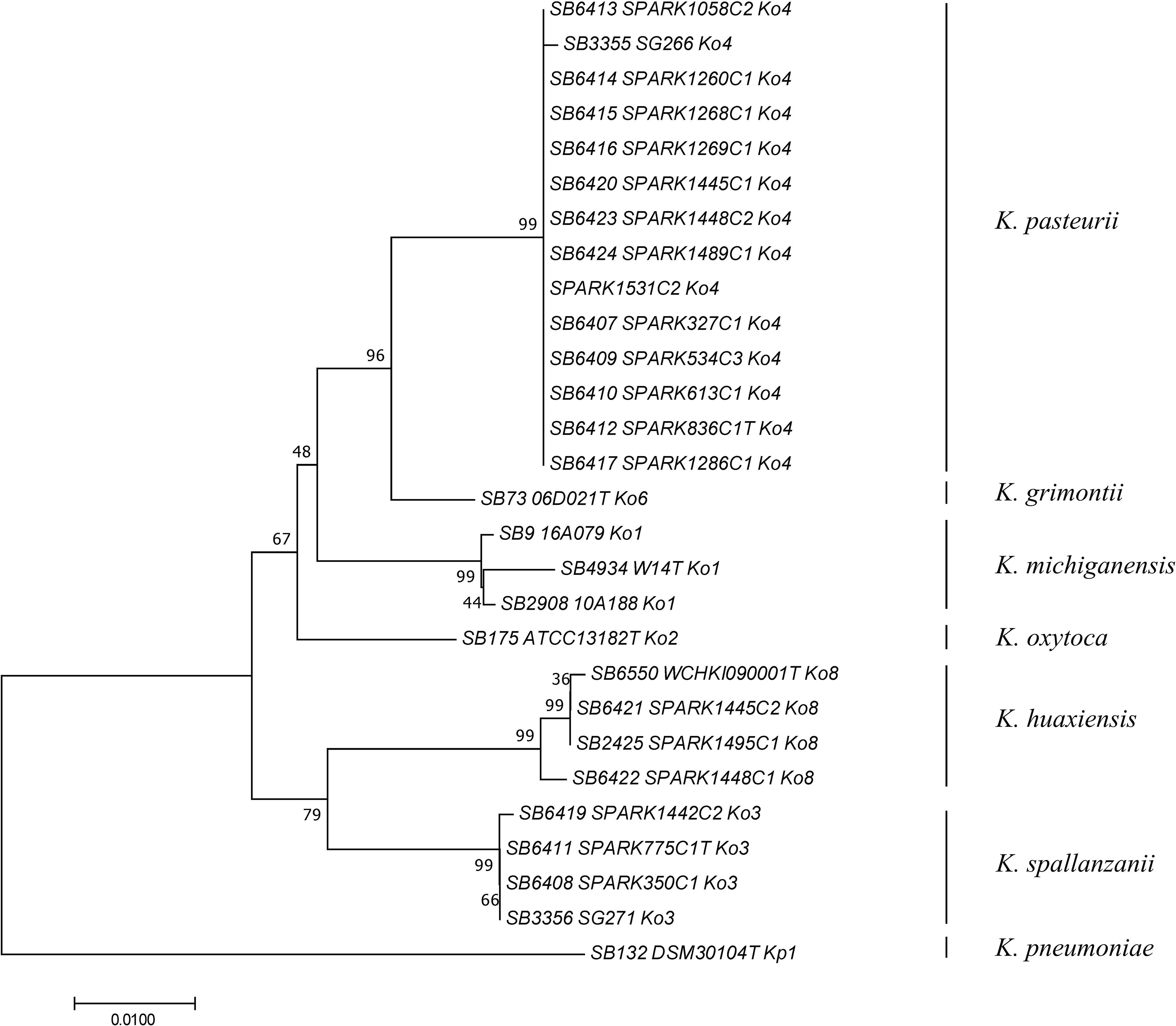
Phylogenetic relationships (neighbor-joining method, Jukes-Cantor correction) based on the concatenated sequences of *gyrA* and *rpoB* genes. The tree was rooted using *K. pneumoniae* DSM 30104^T^ (=ATCC 13883^T^). Taxonomic groups are indicated in front of the branches. Bootstrap proportions obtained after 1000 replicates are indicated at the nodes. Branch lengths represent the number of nucleotide substitutions per site (scale, 0.01 substitution per site). Strain labels are given as Strain Bank ID (e.g., SB73) followed by original strain name, followed by phylogroup. A ‘T’ after the strain name indicates that the strain is the type strain of its taxon.

Average nucleotide identity (ANI) was estimated between Ko3 and Ko4, and the type strains of species of the *K. oxytoca* complex (**Table 2**). The three Ko3 strains, including SPARK_775_C1^T^, shared high identity (above 98%) with the Ko3 strain SG271 (SB3356) (data not shown). The ANI values of SPARK_775_C1^T^ (Ko3) strain with *K. huaxiensis* (WCHKl090001^T^), *K. michiganensis* (W14^T^), *K grimontii* (06D021^T^) and *K. oxytoca* (ATCC 13182^T^) were 90.7%, 88.4%, 88.3% and 87.9%, respectively (**Table 2**). The novel Ko4 strains showed approximately 98% ANI with Ko4 strain SG266 (SB3355). The ANI values of SPARK_836_C1^T^ (Ko4) with *K. grimontii*, *K. michiganensis*, *K. oxytoca* and *K. huaxiensis* were 95.5%, 93.3%, 90.6% and 87.1%, respectively (**Table 2**). Finally, the three Ko8 strains presented ANI values >99% with the type strain of *K. huaxiensis* (WCHKl090001^T^), showing that they belong to this recently described species. The isDDH relatedness range between the Ko3 and Ko4 type strains and other species was 36.3-44.1% and 34.3-67.8%, respectively. In conclusion, both ANI and isDDH values were below the thresholds proposed (30) for species distinction (~95-96% in the case of ANI, ~70% in the case of isDDH), indicating that Ko3 and Ko4 represent two new species.

**Table 2.**
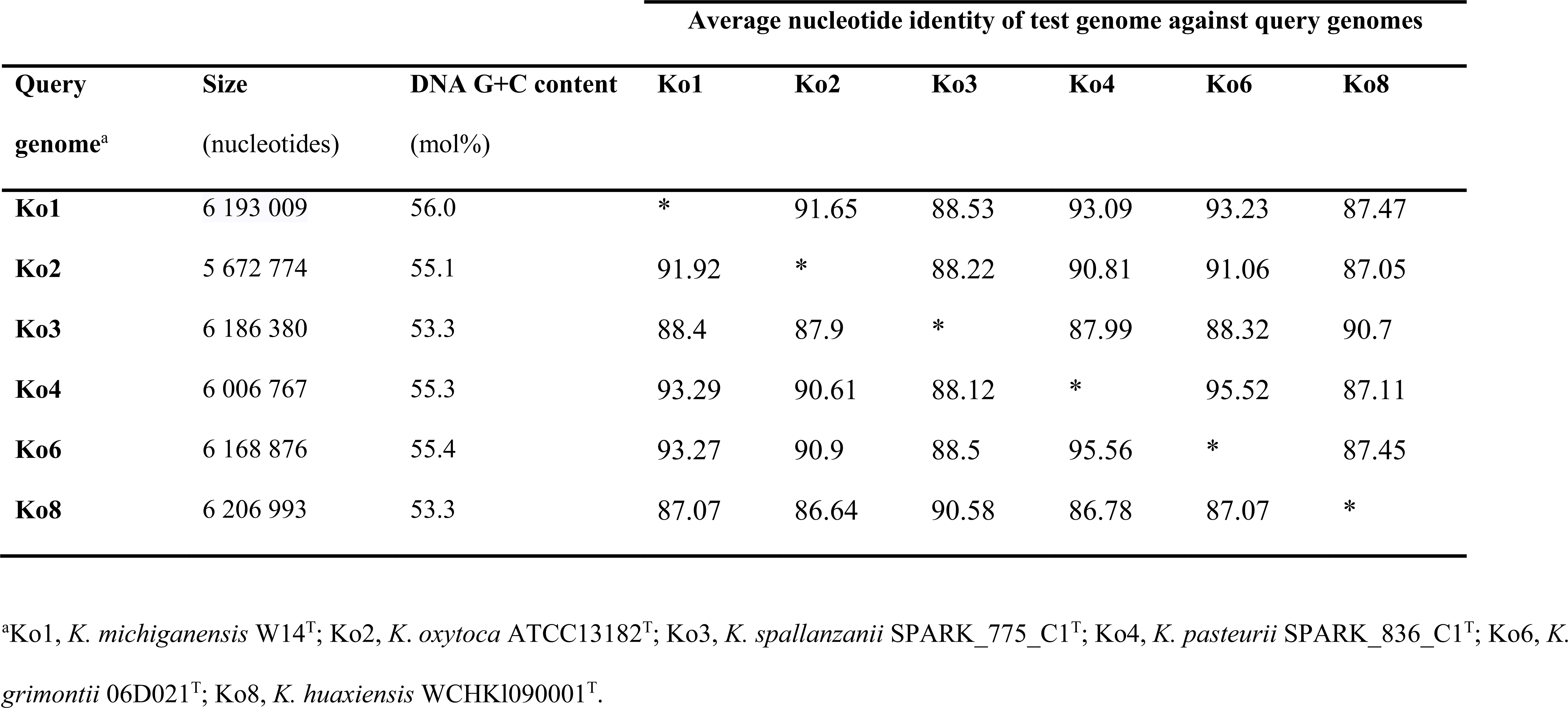
Average nucleotide identity (ANI) values obtained among the type strains of members of the *Klebsiella oxytoca* complex.

The phenotypic characteristics of Ko3 and Ko4 strains were analyzed and compared with those of other *Klebsiella* isolates. We confirmed that all strains were non-motile by microscopy and that all isolates were positive for indole, lactose, mannitol, malonate, lysine decarboxylase and the ONPG test, and reduced nitrate to nitrite, whereas they were all negative for ornithine decarboxylase. Ko8 and Ko3 isolates were negative for Voges–Proskauer test and Ko3 isolates were urease positive (similar to Ko2). To define further the biochemical features of the *K. oxytoca* phylogroups, their carbon source utilization profiles were analysed. Among 190 substrates, several appeared useful for differentiating the phylogroups among themselves and to differentiate Ko3 and Ko4 strains from other groups (**Table 3**, **Figure S5**). The inability to metabolize L-proline and tricarballylic acid differentiated Ko3 strains from other phylogroups except Ko8, which can be differentiated based on its unique ability to utilize 3-O-methyl-glucose. Ko4 had a weak but unique capacity to utilize glyoxylic acid, and differed from Ko6 (*K. grimontii*) by its inability to metabolize D-melezitose; Ko4 was otherwise similar to Ko6 for many features, consistent with their phylogenetic association.

**Table 3.**
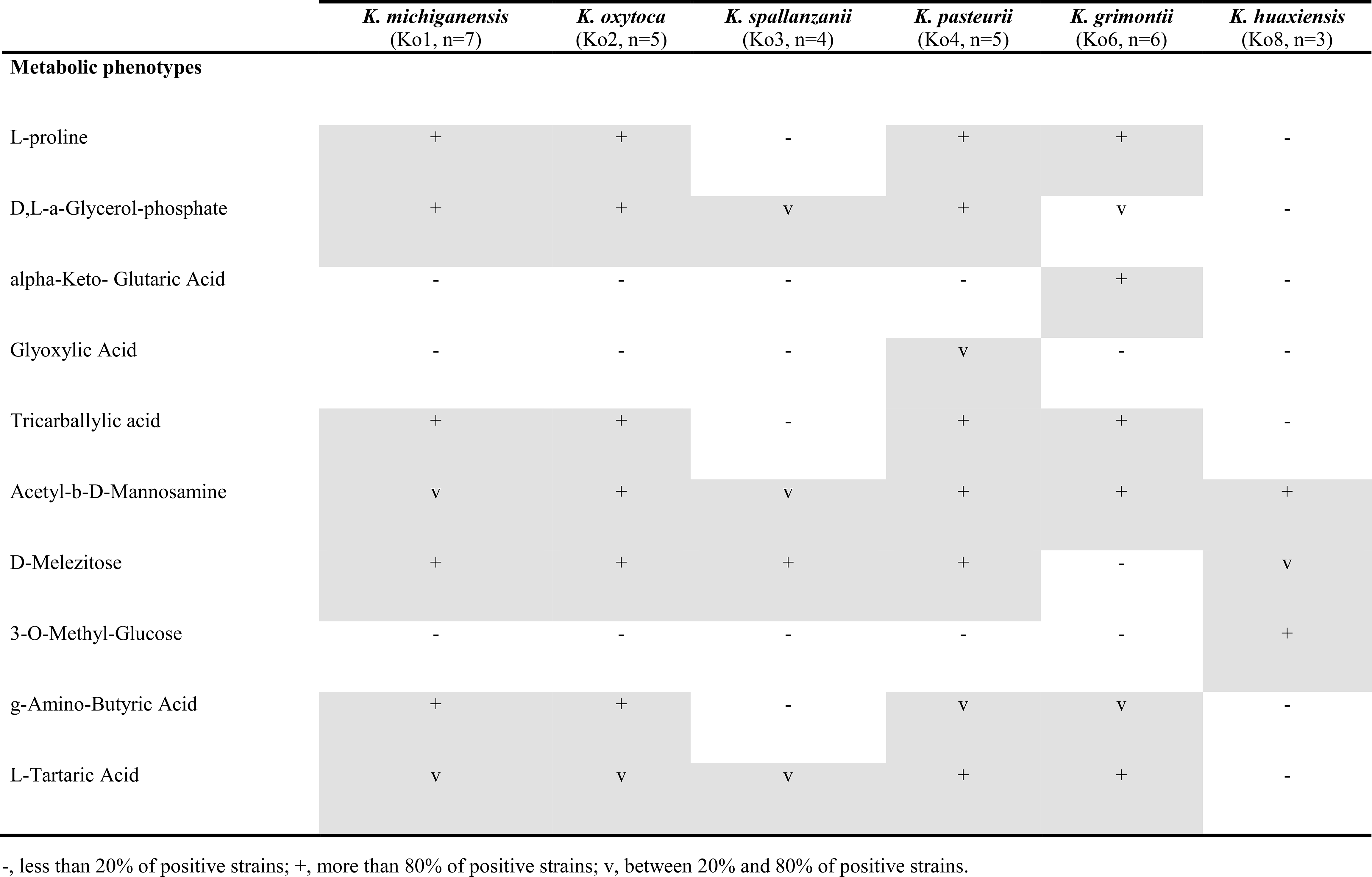
Differential biochemical characteristics of the taxa under study.

We also analysed the MALDI-ToF MS peak patterns of the different members of the *K. oxytoca* complex. Based on the MALDI Biotyper Compass database version 4.1.80 (Bruker Daltonics, Bremen, Germany), the thirty strains were identified either as *K. oxytoca* (23 strains, all belonging to Ko1, Ko2, Ko4, and Ko6) or as *Raoultella ornithinolytica* (7 strains, all strains of Ko3 and Ko8). These misidentifications can be explained by the lack of reference spectra of most phylogroups in the reference database. **Figure S6** summarizes the peak positions found in each strain. A total of 31 biomarkers (2383–10152 *m/z*) associated with specific members of the *K. oxytoca* complex were identified (**Table S1**, **Figure S6**). Consistent with genetic and biochemical findings, we also observed that Ko4 shared most of its spectral peaks with Ko1 and Ko6, presenting only one specific peak (which was variably present) at 3681 *m/z*, whereas Ko3 shared six peaks with only Ko8 and presented two unique peaks at 5178 and 6795 *m/z*. For the remaining phylogroups, specific peaks were observed for Ko2 and Ko8, whereas Ko1 and Ko6 could be identified by specific peak combinations. Based on the current dataset, the specificity and sensitivity of their distribution among phylogroups ranged between 60–100% and 80–100%, respectively (**Table S1**). This finding paves the way to identify isolates of the *K. oxytoca* complex at the species (or phylogroup) level based on MALDI-ToF MS analysis, pending incorporation of reference spectra of the various taxa into reference spectra databases. Based on the above genomic, phenotypic and proteomic characteristics, we propose Ko3 and Ko4 to be considered as two novel species, which we propose to name *K. spallanzanii* and *K. pasteurii*, respectively.

### Description of *Klebsiella spallanzanii* sp. nov

*Klebsiella spallanzanii* (spal. lan.za ‘ni.i N. L. gen. n. referring to Lazzaro Spallanzani, Italian biologist, important contributor to the experimental study of bodily functions and of animal reproduction. He provided what is considered the first disproval of the theory of the spontaneous generation of microbes).

The description is based on 3 strains. Cells are Gram-negative, non-motile, non-spore-forming, straight, rod-shaped and capsulated. Colonies are smooth, circular, white, dome-shaped and glistening. The general characteristics are as described for the genus *Klebsiella*. Indole-positive, ONPG-positive, lysine decarboxylase positive and ornithine decarboxylase negative. Differentiated from the other species of the *K. oxytoca* complex by the urease-positive (similar to Ko2) and Voges–Proskauer test negative (also negative for Ko8). Distinguished from the other members of *K. oxytoca* complex also by the characteristics listed in **Table 3**. Distinguishable from *K. huaxiensis* by the ability to use D-melezitose and the inability to ferment 3-O-methyl-glucose, and from the remaining *K. oxytoca* members by the inability to use L-proline. *K. spallanzanii* isolates were recovered from human urine and cow faeces.

The type strain is strain SPARK_775_C1^T^ (=SB6411, CIP 111695T, DSM 109531), isolated in 2017 from the urine of a patient in Pavia, Italy. The INSDC (GenBank/ENA/DDBJ) accession numbers of the *gyrA*, *rpoB* and *rrs* (coding for 16S rRNA) genes are MN076620, MN076626 and MN091365, respectively. The genome sequence accession number is: *in process*. The DNA G+C content of the type strain is 53.3%.

### Description of *Klebsiella pasteurii* sp. nov

*Klebsiella pasteurii* (pas. teu ‘ri.i N. L. gen. n. referring to Louis Pasteur, a French microbiologist, who made seminal contributions to microbiology and infectious diseases, vaccination and pasteurization. He contributed decisively to disprove the theory of the spontaneous generation of microbes).

The description is based on 14 strains. Cells are Gram-negative, non-motile, non-spore-forming, straight, rod-shaped and capsulated. Colonies are smooth, circular, white, dome-shaped and glistening. The general characteristics are as described for the genus *Klebsiella*. Indole-positive, urease-negative, ONPG-positive, Voges–Proskauer test positive, lysine decarboxylase positive and ornithine decarboxylase negative. Distinguished from the other members of *K. oxytoca* complex by the characteristics listed in **Table 3**. Distinguishable from *K. grimontii* by the ability to ferment D-melezitose and inability to ferment alpha-keto-glutaric acid, and from the remaining *K. oxytoca* groups by the unique weak ability to ferment glyoxylic acid. *K. pasteurii* isolates were recovered from faeces of cows, turtles and humans.

The type strain is strain SPARK_836_C1^T^ (=SB6412, CIP 111696T, DSM 109530), isolated in 2017 from the faeces of a patient in Pavia, Italy. The INSDC (GenBank/ENA/DDBJ) accession numbers of the *gyrA*, *rpoB* and *rrs* (coding for 16S rRNA) genes are MN076619, MN076625 and MN091366, respectively. The genome sequence accession number is: *in process*. The DNA G+C content of the type strain is 55.4%.

## Author contributions

Isolation of *Klebsiella* from diverse sources: CM, MC, PM, CB, DS; Microbiological characterization of isolates: CM, CR, VP, MC. Genomic sequencing: CM, HAT, TVSK, DS. Sequence data analysis: CM, CR, HAT, TVSK. MALDI-TOF analysis: CR, VP. Phenotypic microarray analyses: VP, SB. Initial manuscript writing: CM, CR, SB. Manuscript revision: all. Funding acquisition: EJF, SB, JC, CB, DS.

## Acknowledgements

We acknowledge Marie-Hélène Nicolas-Chanoine and Alan McNally for providing strains included in this study.

## References

1. Brisse S, Grimont F, Grimont PAD. “The genus Klebsiella,”. In Dworkin M, Falkow S, Rosenberg E, Schleifer K-H, Stackebrandt E, editors. The Prokaryotes-A Handbook on the Biology of Bacteria. New York, NY: Springer (2006). p. 159–197.

2. Caltagirone M, Nucleo E, Spalla M, Zara F, Novazzi F, Marchetti VM, et al. Occurrence of Extended Spectrum β-Lactamases, KPC-Type, and MCR-1.2-Producing Enterobacteriaceae from Wells, River Water, and Wastewater Treatment Plants in Oltrepò Pavese Area, Northern Italy. Front microbiol (2017) 8:2232. 10.3389/fmicb.2017.02232.

3. Schmitz RA, Klopprogge K, Grabbe R. Regulation of nitrogen fixation in Klebsiella pneumoniae and Azotobacter vinelandii: NifL, transducing two environmental signals to the nif transcriptional activator NifA. J Mol Microbiol Biotechnol (2002) 4(3):235–42.

4. Paczosa MK, Mecsas J. Klebsiella pneumoniae: Going on the Offense with a Strong Defense. Microbiol Mol Biol Rev (2018) 80(3):629–661. 10.1128/MMBR.00078-15.

5. Broberg CA, Palacios M, Miller VL. Klebsiella: a long way to go towards understanding this enigmatic jet-setter. F1000 Prime Rep (2014) 6:64. 10.12703/P6-64.

6. Fournier B, Roy PH. Variability of Chromosomally Encoded b-Lactamases from Klebsiella oxytoca. Antimicrob Agents Chemother (1997) 41(8):1641–1648.

7. Granier SA, Plaisance L, Leflon-Guibout V, Lagier E, Morand S, Goldstein FW, et al. Recognition of two genetic groups in the Klebsiella oxytoca taxon on the basis of chromosomal b-lactamase and housekeeping gene sequences as well as ERIC-1 R PCR typing. Int J Syst Evol Microbiol (2003) 53:661–668.10.1099/ijs.0.02408-0.

8. Granier SA, Leflon-Guibout V, Goldstein FW, Nicolas-Chanoine MH. New Klebsiella oxytoca beta-lactamase genes bla(OXY-3) and bla(OXY-4) and a third genetic group of K oxytoca based on bla (OXY-3). Antimicrob Agents Chemother (2003) 47:2922–2928. 10.1128/AAC.47.9.2922-2928.2003.

9. Fevre C, Jbel M, Passet V, Weill FX, Grimont PA, Brisse S. Six groups of the OXY b-lactamase evolved over millions of years in Klebsiella oxytoca. Antimicrob Agents Chemother (2005) 49:3453–3462. 10.1128/AAC.49.8.3453-3462.2005.

10. Izdebski R, Fiett J, Urbanowicz P, Baraniak A, Derde LP, Bonten MJ, et al. Phylogenetic lineages, clones and β-lactamases in an international collection of Klebsiella oxytoca isolates non susceptible to expanded-spectrum cephalosporins. J Antimicrob Chemother (2015) 70(12):3230–7. 10.1093/jac/dkv273.

11. Saha R, Farrance CE, Verghese B, Hong S, Donofrio RS. Klebsiella michiganensis sp. nov., a new bacterium isolated from a toothbrush holder. Curr Microbiol (2013) 66:72–78. 10.1007/s00284-012-0245-x.

12. Passet V, Brisse S. Description of Klebsiella grimontii sp. nov. Int J Syst Evol Microbiol (2018) 68:377–381. 10.1099/ijsem.0.00251

13. Hu Y, Wei L, FengY, Xie Y, Zong Z. Klebsiella huaxiensis sp. nov., recovered from human urine. Int J Syst Evol Microbiol (2019) 69(2):333–336. 10.1099/ijsem.0.003102.

14. Van Kregten E, Westerdaal NA, Willers JM. New, simple medium for selective recovery of Klebsiella pneumoniae and Klebsiella oxytoca from human feces. J Clin Microbiol (1984) 20(5):936–41.

15. Brisse S, Passet V, Grimont PA. Description of Klebsiella quasipneumoniae sp. nov., isolated from human infections, with two subspecies, Klebsiella quasipneumoniae subsp. Quasipneumoniae subsp. nov. and Klebsiella quasipneumoniae subsp. similipneumoniae subsp. nov., and demonstration that Klebsiella singaporensis is a junior heterotypic synonym of Klebsiella variicola. Int J Syst Evol Microbiol (2014) 64:3146–3152. 10.1099/ijs.0.062737-0.

16. Seemann T. Prokka: rapid prokaryotic genome annotation. Bioinformatics (2014) 30(14):2068–9. 10.1093/bioinformatics/btu153.

17. Richter M, Rosselló-Móra R, Glöckner FO, Peplies J. JSpeciesWS: a web server for prokaryotic species circumscription based on pairwise genome comparison. Bioinformatics (2016) 32(6):929–31. 10.1093/bioinformatics/btv681.

18. Meier-Kolthoff JP, Auch AF, Klenk HP, Göker M. Genome sequence-based species delimitation with confidence intervals and improved distance functions. BMC Bioinformatics (2013) 14:60.

19. Edgar RC. MUSCLE: multiple sequence alignment with high accuracy and high throughput Nucleic Acids Res (2004) 32(5):1792–1797. 10.1093/nar/gkh340.

20. Kumar S, Stecher G, Tamura K. MEGA7: Molecular Evolutionary Genetics Analysis Version 7.0 for Bigger Datasets. Mol Biol Evol (2016) 33(7):1870–4. 10.1093/molbev/msw054.

21. Jukes TH, Cantor CR. “Evolution of protein molecules”. In: Munro HN, editor. Mammalian Protein Metabolism, New York, NY: Academic Press (1969) p. 21–132.

22. Jones DT, Taylor WR, Thornton JM. The rapid generation of mutation data matrices from protein sequences. Comput Appl Biosci (1992) 8:275–282.

23. Page AJ, Cummins CA, Hunt M, Wong VK, Reuter S, Holden MTG, et al. Roary: Rapid large-scale prokaryote pan genome analysis. Bioinformatics (2015). 31(22):3691–3693. 10.1093/bioinformatics/btv421.

24. Price MN, Dehal PS, Arkin AP. FastTree 2 - Approximately Maximum-Likelihood Trees for Large Alignments. PLoS ONE (2010) 5(3):e9490. 10.1371/journal.pone.0009490.

25. Blin C, Passet V, Touchon M, Rocha EPC, Brisse S. Metabolic diversity of the emerging pathogenic lineages of Klebsiella pneumoniae. Environ Microbiol (2017) 19(5):1881–1898. 10.1111/1462-2920.13689.

26. Rodrigues C, Passet V, Rakotondrasoa A, Brisse S. Identification of Klebsiella pneumoniae, Klebsiella quasipneumoniae, Klebsiella variicola and Related Phylogroups by MALDI-TOF Mass Spectrometry. Front Microbiol (2018) 9:3000. 10.3389/fmicb.2018.03000.

27. Brisse S, Verhoef J. Phylogenetic diversity of Klebsiella pneumoniae and Klebsiella oxytoca clinical isolates revealed by randomly amplified polymorphic DNA, gyrA and parC genes sequencing and automated ribotyping. Int J Syst Evol Microbiol (2001) 51:915–24. 10.1099/00207713-51-3-915.

28. Naum M, Brown EW, Mason-Gamer RJ. Is 16S rDNA a reliable phylogenetic marker to characterize relationships below the family level in the Enterobacteriaceae? J Mol Evol (2008) 66(6):630–42. 10.1007/s00239-008-9115-3.

29. Boye K, Hansen DS. Sequencing of 16S rDNA of Klebsiella: taxonomic relations within the genus and to other Enterobacteriaceae. Int J Med Microbiol (2003) 292(7–8):495–503. 10.1078/1438-4221-00228.

30. Rossello-Mora R, Amann R. Past and future species definitions for Bacteria and Archaea. Syst Appl Microbiol (2015) 38:209–216. 10.1016/j.syapm.2015.02.001.

