## Supplementary figures and images for "Description of *Klebsiella spallanzanii* sp. nov. and of *Klebsiella pasteurii* sp. nov"

### Suppl Figure S1

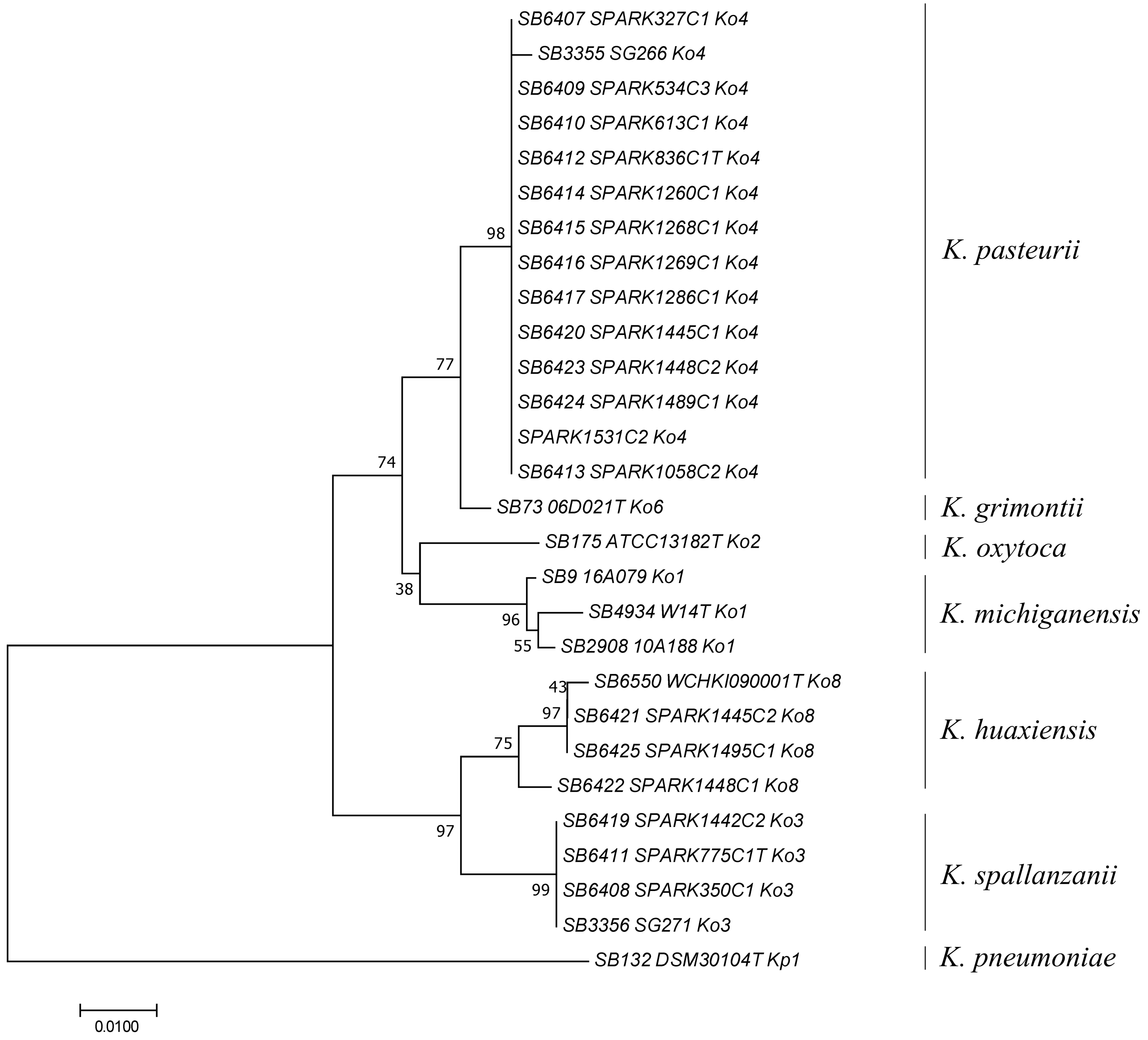

### Suppl Figure S2

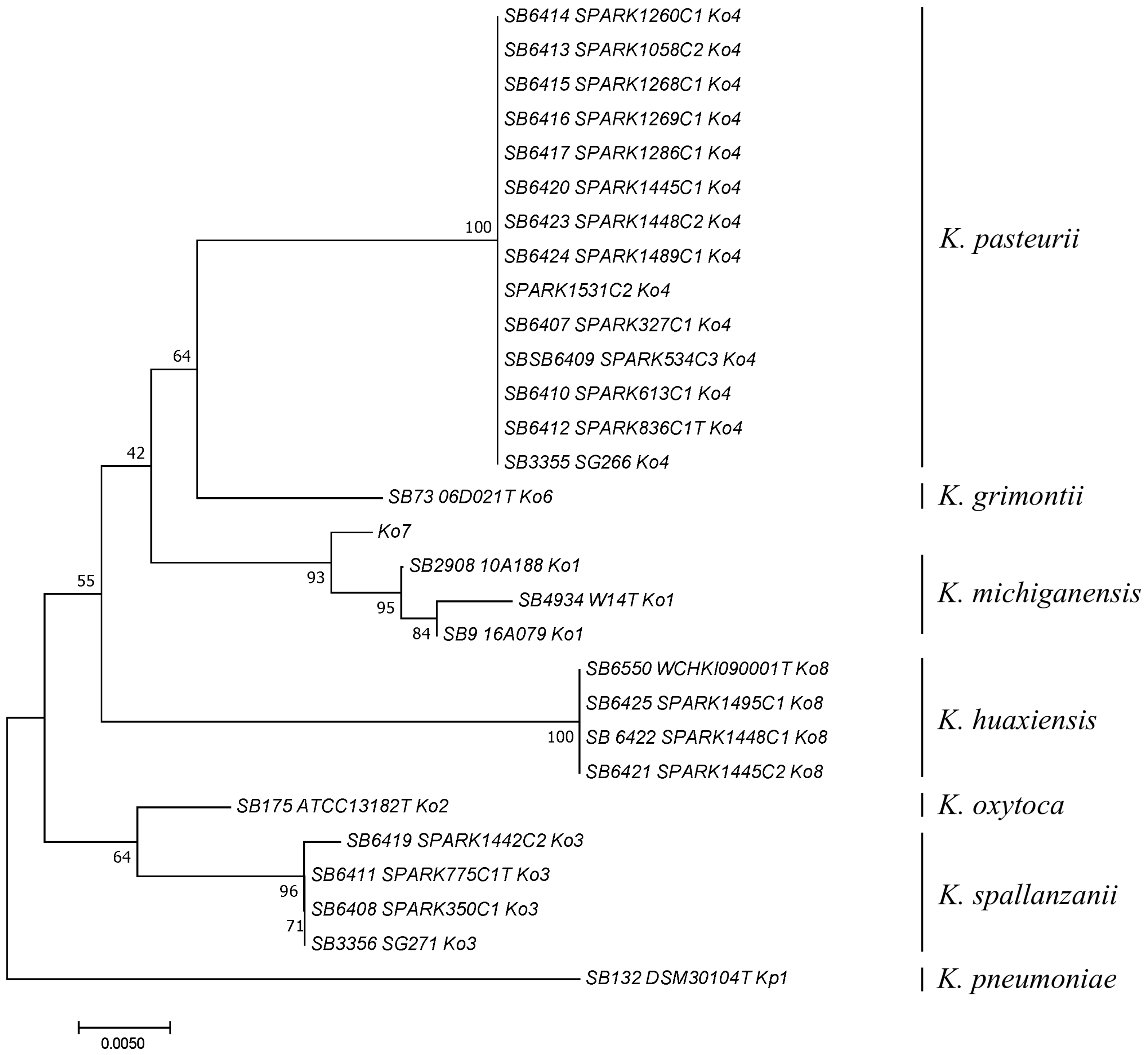

### Suppl Figure S3

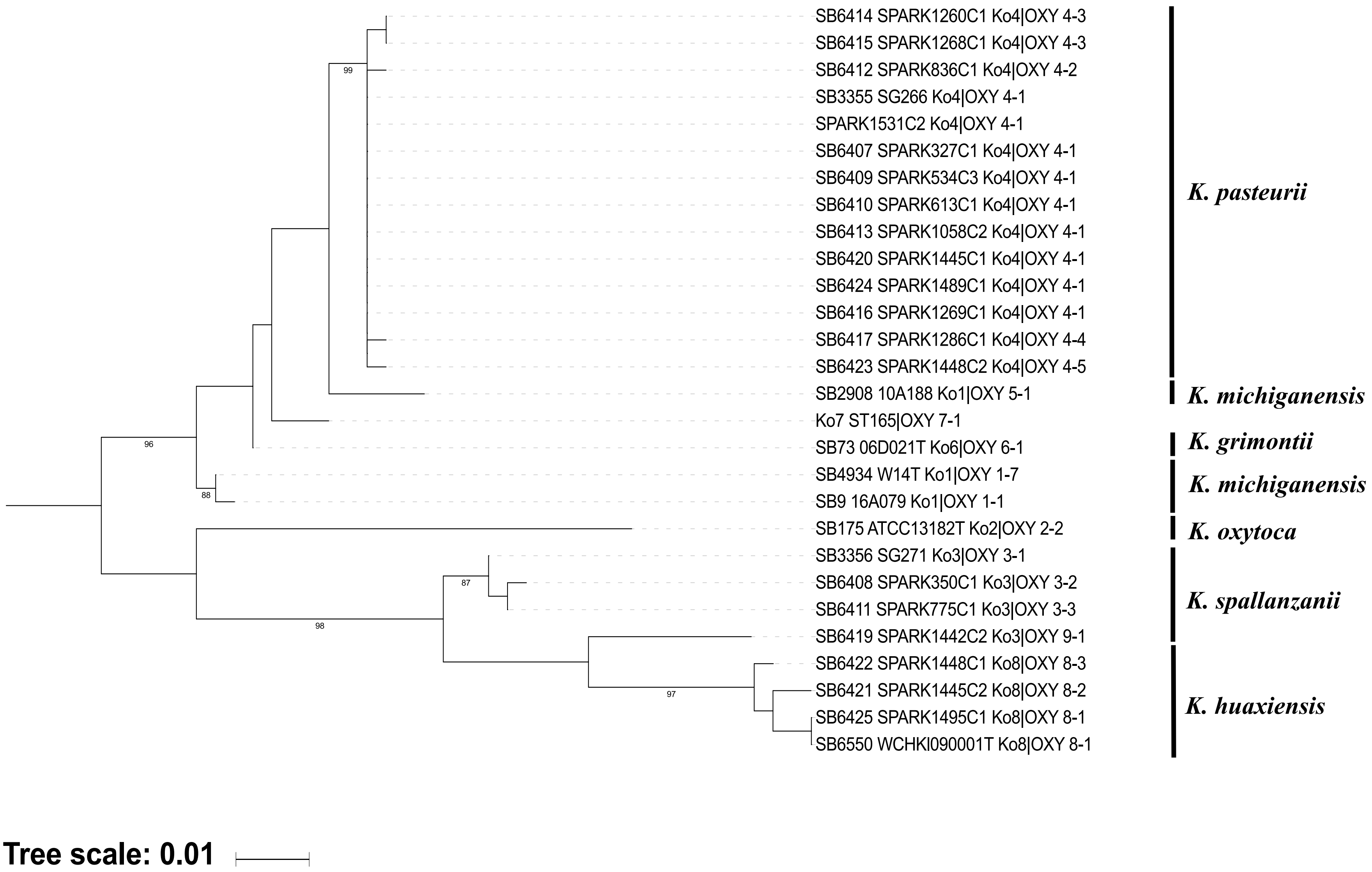

### Suppl Figure S4

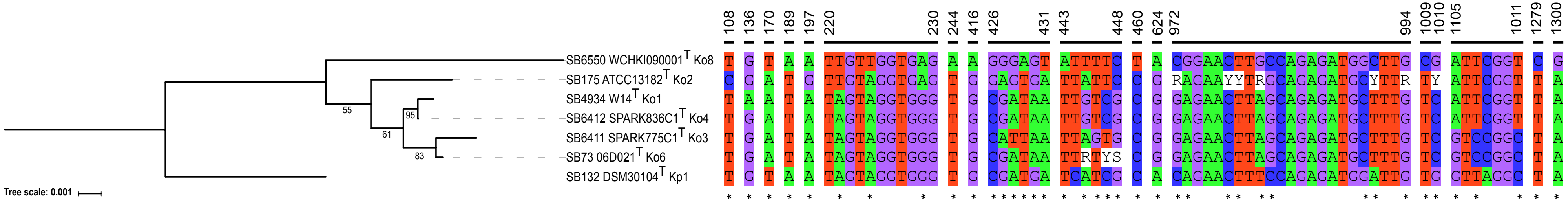

### Suppl Figure S6

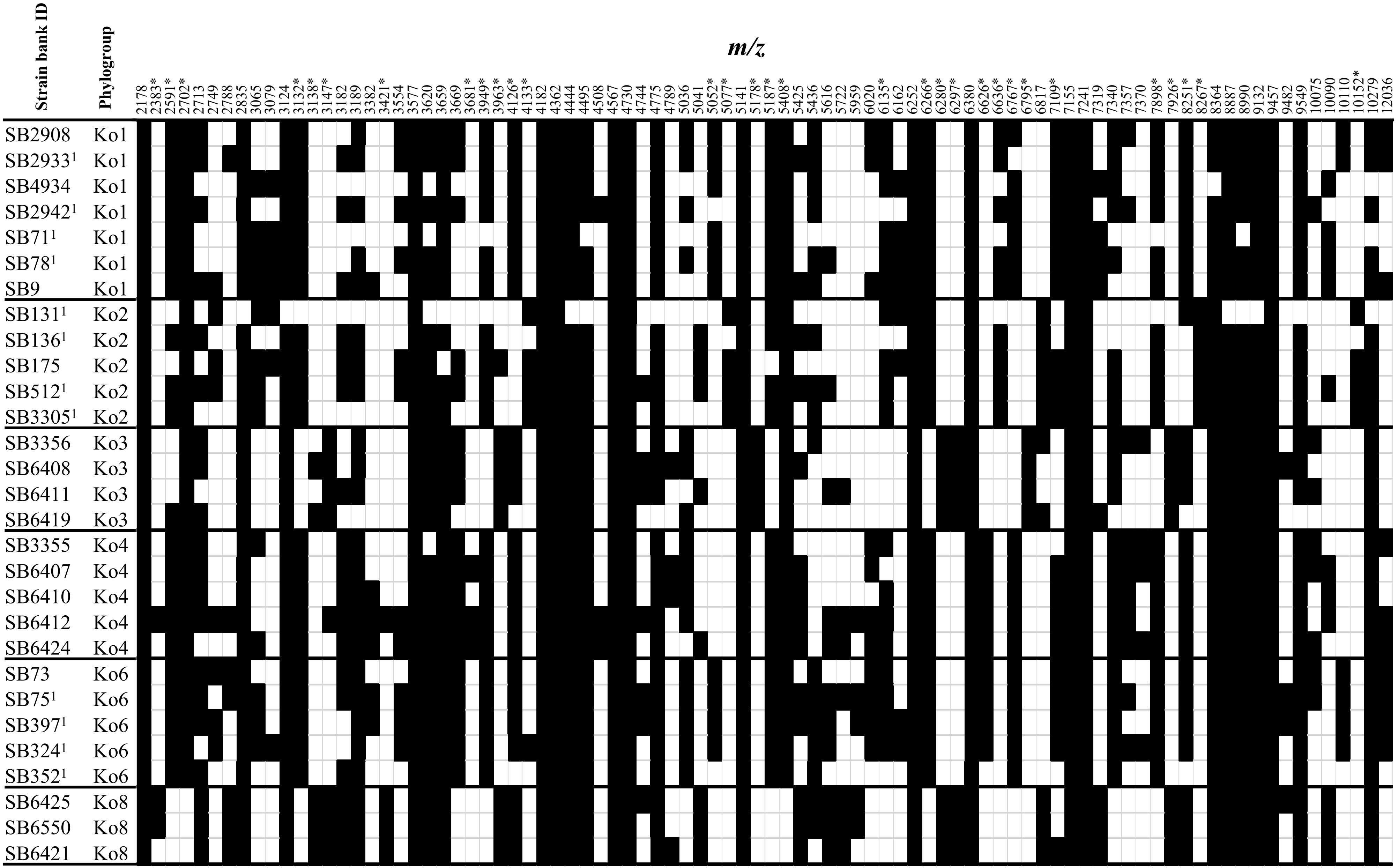
