## Supplementary material for "Description of *Klebsiella spallanzanii* sp. nov. and of *Klebsiella pasteurii* sp. nov": Suppl Figure S5

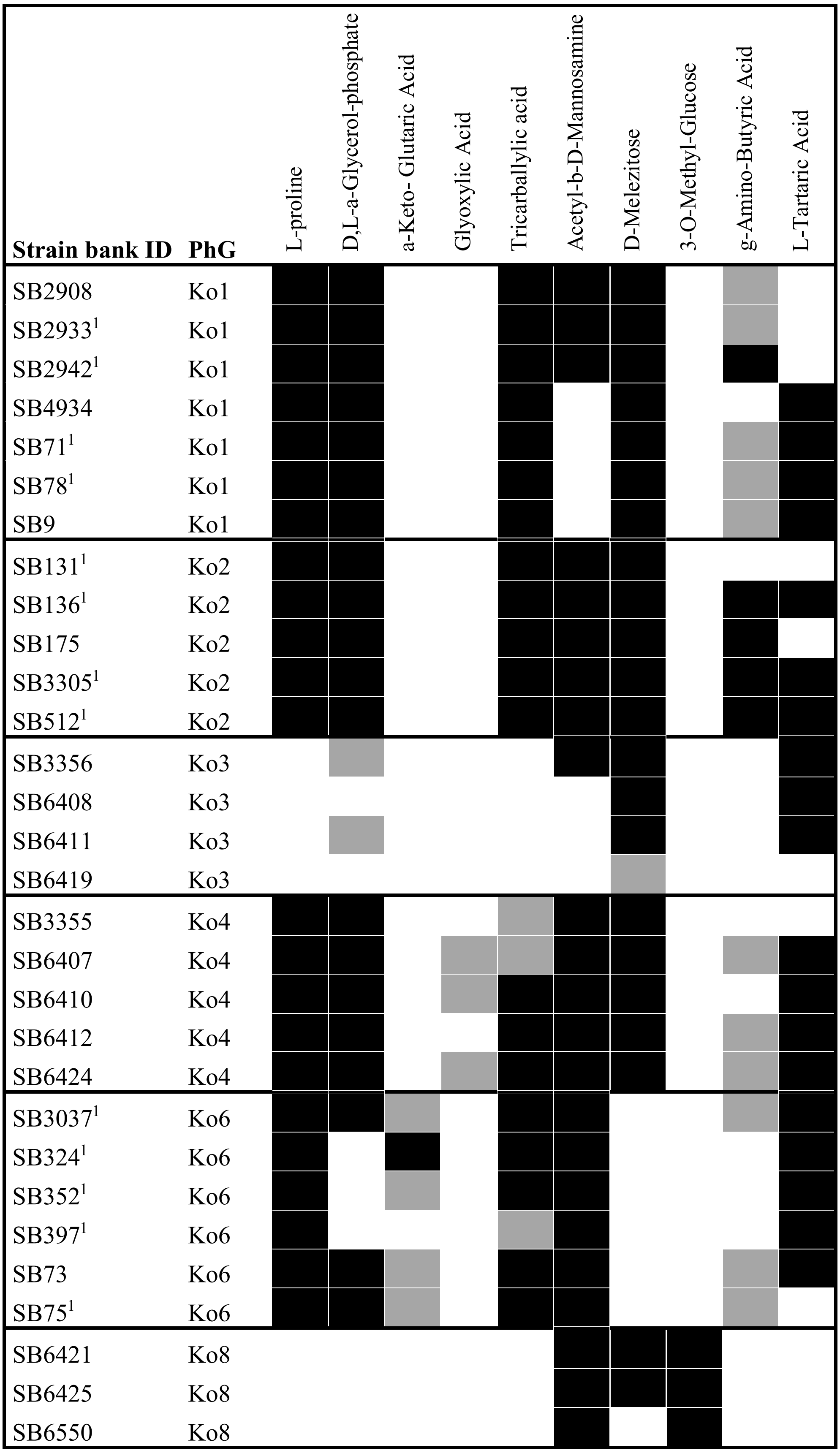


PhG, Phylogroup; [1]Strains added to the study for phenotype microarray experiments (Biolog)
