## Supplementary material for "Description of *Klebsiella spallanzanii* sp. nov. and of *Klebsiella pasteurii* sp. nov": Table S1

**Table S1**. MALDI-ToF mass spectrometry peaks, which are useful biomarkers to discriminate phylogroups of the *Klebsiella oxytoca* species complex.

| Phylogroup(s) in which the peak was observed | Peak Position (*m/z*)[1] | Sensitivity [95% CI] | Specificity [95% CI] |
|---|---|---|---|
| **Ko1, Ko4, Ko6, Ko3, Ko2** | 2702[2] | 100% [86.77% - 100.00%] | 100% [29.24% - 100.00%] |
| | 5408 | 100% [86.77% - 100.00%] | 100% [29.24% - 100.00%] |
| **Ko1, Ko4, Ko6, Ko3, Ko8** | 4126[2] | 91.67% [73.00% - 98.97%] | 100% [47.82% - 100.00%] |
| | 8251 | 91.67% [73.00% - 98.97%] | 80% [28.36% - 99.49%] |
| **Ko1, Ko4, Ko6, Ko2** | 2591[2] | 90.91% [70.84% - 98.88%] | 85.71% [42.13% - 99.64%] |
| | 5187 | 95.45% [77.16% - 99.88%] | 100% [59.04% - 100%] |
| | 3132[2] | 95.45% [77.16% - 99.88%] | 100% [59.04% - 100%] |
| | 6266 | 100% [84.56% - 100%] | 100% [59.04% - 100%] |
| | 3949[2] | 90.91% [70.84% - 98.88%] | 100% [59.04% - 100%] |
| | 7898 | 90.91% [70.84% - 98.88%] | 100% [59.04% - 100%] |
| | 6135 | 81.82% [59.72% - 94.81%] | 100% [59.04% - 100%] |
| | 7109 | 81.82% [59.72% - 94.81%] | 85.71% [42.13% - 99.64%] |
| | 6767 | 94.12% [71.31% - 99.85%] | 100% [73.54% - 100%] |
| **Ko1, Ko6** | 5052 | 83.33% [51.59% - 97.91%] | 100% [80.49% - 100.00%] |
| **Ko6, Ko4** | 6626 | 90% [55.50% - 99.75%] | 100% [82.35% - 100.00%] |
| **Ko4** | 3681 | 60% [14.66% - 94.73%] | 100% [85.75% - 100%] |
| **Ko1, Ko2** | 6636 | 85.71% [42.13% - 99.64%] | 100% [78.20% - 100%] |
| **Ko2** | 4133[2] | 80% [28.36% - 99.49%] | 95.83% [78.88% - 99.88%] |
| | 8267 | 100% [47.82% - 100.00%] | 100% [85.75% - 100.00%] |
| | 5077[2] | 80% [28.36% - 99.49%] | 100% [85.75% - 100.00%] |
| | 10152 | 80% [28.36% - 99.49%] | 100% [85.75% - 100.00%] |
| **Ko3, Ko8** | 3138[2] | 71.43% [29.04% - 96.33%] | 100% [84.56% - 100%] |
| | 6280 | 100% [59.04% - 100.00%] | 100% [84.56% - 100%] |
| | 3147[2] | 100% [59.04% - 100.00%] | 95.45% [77.16% - 99.88%] |
| | 6297 | 100% [59.04% - 100.00%] | 100% [84.56% - 100%] |
| | 3963[2] | 100% [59.04% - 100.00%] | 96.15% [80.36% - 99.90%] |
| | 7926 | 100% [59.04% - 100.00%] | 100% [84.56% - 100%] |
| **Ko3** | 5178 | 100% [39.76% - 100.00%] | 100% [86.28% - 100%] |
| | 6795 | 100% [39.76% - 100.00%] | 100% [86.28% - 100%] |
| **Ko8** | 2383 | 66.67% [9.43% - 99.16%] | 96.15% [80.36% - 99.90%] |
| | 3421 | 100% [29.24% - 100.00%] | 96.15% [80.36% - 99.90%] |

CI, confidence interval
[1] Position in the spectra using a tolerance of ±0.03%.
[2] Double-charged ion.
